# Versatile HIV Rev-dependent reporter cell system for stringent and sensitive quantification of viral reservoirs, neutralizing antibodies, and restriction factors

**DOI:** 10.1101/2025.09.08.674319

**Authors:** Mark Spear, Joseph Choi, Brian Hetrick, JeongHwan Lee, Gabrielle Lê-Bury, Thuy T. Vo, Yang Han, Huizhi Liang, Jia Guo, Dongyang Yu, Subashini Iyer, Henry C. Mwandumba, David G. Russell, Yuntao Wu, David W. Gludish

## Abstract

Detecting and measuring HIV reservoirs, neutralizing antibodies, and restriction factors are important for HIV cure research and the development of new therapeutics and vaccines. Here we describe the development and validation of several HIV Rev-dependent indicator cell lines for these purposes. These reporter cells come from different T-lymphoblast cell lines, including MOLT-4-R5, SupT1-R5, CEM-SS, A3R5, and from the adherent TZM cell platform based on HeLa clone JC53. These cells express CD4, CXCR4, and various levels of CCR5. We compared these cell lines for responsiveness to both X4 and R5-tropic viruses, and confirmed that reporter expression in these cells is not affected by stimulation from mitogens but is responsive to HIV Tat and Rev, reducing non-specific reporter induction from the leaky LTR promoter. To validate the sensitivity of the Rev-dependent reporter cell systems, we conducted a viral dilution assay with three primary HIV-1 clade C swarms from an adult in Malawi. We also validated the systems for quantifying antibody neutralization and screening restriction factors; these systems are also sensitive for viral outgrowth assays for quantifying viral reservoirs in clinical and basic research settings. Given that the systems can measure HIV accurately in complex environments with mitogens or other substances, they can be used for versatile applications, such as quantifying latent reservoirs, testing inhibitory compounds, conducting neutralizing antibody assays, and identifying new restriction factors.

## Introduction

The development of facile and sensitive reporter cell systems is critical to HIV research, enabling routine quantification of viral titer, replication kinetics, and inhibition of infection while also allowing characterization of patient viral reservoirs. The first such systems employed the HIV long terminal repeat (LTR) to drive reporter gene expression [1-5]. In these systems, Tat upregulates gene expression from the LTR in at least two non-exclusive mechanisms: (1) co-transcriptional anti-termination through binding to the trans-activation response (TAR) element in nascent transcripts, increasing elongation efficiency [6-11]; and (2) recruitment of transcription factors to the RNA Polymerase II pre-initiation complex at the LTR, promoting transcription initiation [12, 13]. However, Tat-independent expression from the LTR must also occur early during infection to permit Tat accumulation and subsequent Tat-dependent expression: the proviral LTR and associated reporter systems are prone to Tat-independent induction of gene expression by mitogens, cytokines, and other stimuli [14-16].This leakiness can confound endpoints and hinder reporter implementation in high-sensitivity applications, like viral outgrowth assays (VOAs), or where LTR-modulating compounds may be present, such as screens for neutralizing antibodies in plasma or for small molecule inhibitors.

To redress this, we previously [17] developed the Rev-dependent HIV reporter vector technology. These systems retain the viral LTR and its partial Tat dependence, but implement an additional layer of post-transcriptional regulation mediated by HIV Rev. In Rev-dependent vectors, the reporter cDNA is juxtaposed with the Rev-response element (RRE) and interposed between a viral splice donor and acceptor [17, 18]. In the absence of Rev protein, unspliced transcripts are retained in the nucleus until splicing is complete, eliminating the reporter mRNA in the process. During infection, intracellular Rev derived from HIV proviral transcription binds to unspliced reporter transcripts and promotes their nucleocytoplasmic export, upregulating expression [17, 19-22]. This Boolean AND requirement for intracellular Tat and Rev substantially increases the specificity of reporter expression [23]. The reduction in HIV-independent background reporter expression afforded by these systems renders them singularly well-suited for applications requiring high specificity and sensitivity. In one such application, Rev-dependent technology was employed for targeted HIV and SIV reservoir depletion through expression of cytotoxic proteins [24, 25]. Additionally, these vectors have been used to generate several Rev-dependent reporter cell lines, including Affinofile-293 [26], Rev-CEM [18], TZM-gfp [23], and Sup-GGR [27].

Here, we report the development of additional and novel Rev-dependent reporter cells: derived from the T -lymphoblast cell lines, MOLT-4-R5, SupT1-R5, and A3R5, which are CCR5-expressing derivatives of MOLT-4 [23], SUP-T1 [28], and CEM A3.01 [29]; derived from the CCR5-/Lo line, CEM-SS [30], a syncytium-sensitive CEM derivative; and several additional reporter variants of the TZM platform derived from HeLa cell clone JC53 [5]. While each of these cell lines exhibited robust and specific detection of HIV, they demonstrated variable responsiveness to X4 and R5-tropic HIV-1NL4-3 and HIV-1JR-CSF, congruent with anticipated clonal variation. We confirmed reporter gene Rev-dependency using a ΔRev HIV-1 NL4-3 mutant [31] and with a Tat and Rev-expressing lentivirus. We also demonstrate recalcitrance to mitogen-induced reporter expression, arguing that the requirement for both intracellular Tat and Rev suppresses spurious expression from leaky LTR expression. To demonstrate the high sensitivity of the Rev-dependent reporter cell system to clinical isolates, we performed a VOA and cell-free viral dilution assay comparing three primary HIV-1 clade C viral swarms isolated from a single Malawian adult. Lastly, to highlight other relevant use cases for these systems, antibody neutralization and restriction factor screening assays were performed.

The Rev-dependent reporter cell lines described here constitute a versatile toolkit in HIV research. These systems are sufficiently sensitive for VOAs, can supplement or replace conventional p24 ELISAs in clinical and basic research, and exhibit the required specificity to measure HIV output in complex matrices containing mitogens or other perturbagens. These systems are well suited for a range of end uses including latent reservoir quantification by VOA, inhibitory compound screens, neutralizing antibody assays, and the identification of novel restriction factors.

## Results

### Development of multiple Rev-dependent reporter cells for divergent applications in HIV research

The original HIV-1 Rev-dependent lentiviral vector was developed by Wu et al. [17] to identify and target HIV-positive cells [24, 25, 32]. The vector was subsequently used to establish an HIV Rev-dependent reporter T cell line, Rev-CEM [18], which enables the quantification of low-level Tat and Rev expression from the unintegrated HIV genome [33]. Accurately measuring low-level HIV gene expression poses challenges with common Tat-dependent, LTR-driven reporters, primarily because leaky LTR transcription can obscure true Tat responses [18]. Tat-independent transactivation of the LTR is well documented to occur through stimulation with cytokines, mitogens, transcription activators, HDAC inhibitors, or simply due to HIV structural proteins such as Env on the virion [15, 16, 34-42]. The Rev-dependent vector introduces additional regulation via the Rev-responsive element (RRE) and multiple authentic splicing sites from the HIV genome to enhance reporter specificity and HIV dependence (**Fig. 1A**). While the LTR responds to both Tat and non-viral stimuli, placing the reporter cDNA as an intron upstream of the RRE adds a requirement of Rev protein expression for the nuclear export and translation of unspliced reporter mRNAs [17, 21].

**Fig. 1.**
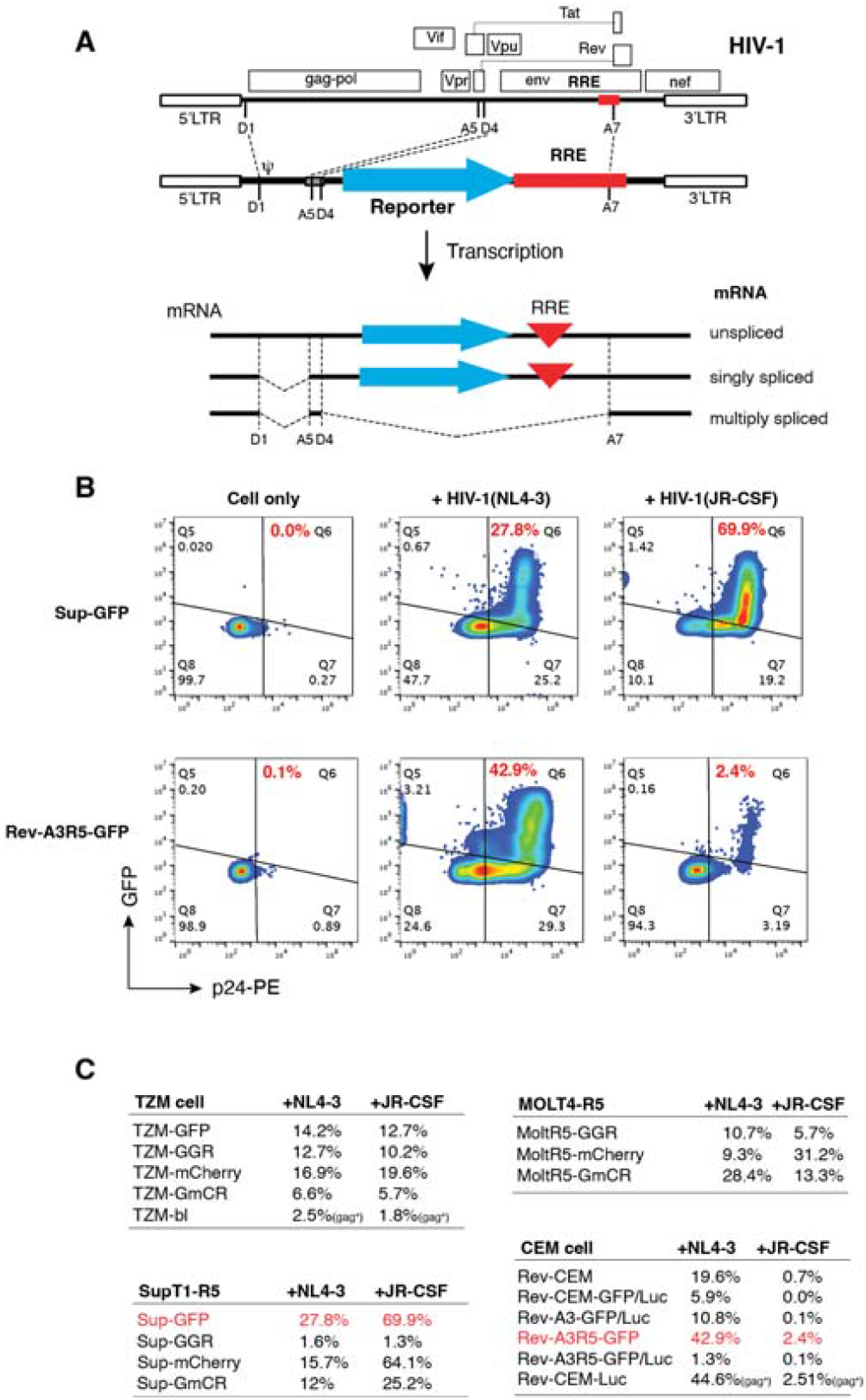
Development and comparison of multiple Rev-dependent reporter cells. **(A)** Illustration of the Rev-dependent lentiviral vector and its transcription. The vector is based on the genome of HIV-1(NL4-3). The reporter gene is inserted upstream of the RRE and between the HIV splice donor (D4) and the splice acceptor (A7). Transcription starts in the upstream LTR and is stimulated by the Tat-mediated transactivation. Complete mRNA splicing removes the reporter open reading frame (ORF) and RRE, whereas partial and complete splicing retains the reporter ORF and RRE that are dependent on Rev for nuclear export and translation. (**B** and **C**) Comparison of reporter responsiveness of the reporter cells listed in (**C)** to HIV-1(NL4-3) and HIV-1(JR-CSF). Cells were identically infected with HIV-1(NL4-3) or HIV-1(JR-CSF) targeting MOI 0.1 (TZM-GFP) for 2 hours, and then washed, and cultured for 2 days.

The Rev-CEM T cell line has been utilized in various laboratories for research on HIV cell-cell transmission, screening of Rev inhibitors, and studying antiviral drug resistance [33, 43-49]. However, the cell has limitations, including a lack of versatility in reporters and the ability to quantify M-tropic viruses, which has prompted its re-cloning and reengineering for applications like viral tropism typing [26, 49]. In addition, we and others have created more versatile reporter cell lines, including Sup-GGR [27], TZM-gfp [23], and Rev-A3R5-GFP [50], Affinofile 293-GGR [26] for viral outgrowth assays and the analysis of early HIV-1 infections [23].

We developed and compared several cell lines for their specificity and performance in reporting infection with either X4-tropic (NL4-3) and R5-tropic (JR-CSF) HIV-1 (**Fig. 1B** and **1C**). For comparison, the Tat-dependent cell line TZM-bl was also included [5, 51]. These cells were infected identically (with an equal level of p24 input) and analyzed for reporter expression using flow cytometry. We also conducted intracellular staining of HIV p24 to correlate HIV-positive cells with reporter expression (**Fig. S1**). Representative results are shown in **Fig. 1B**. In uninfected cells, we detected minimal p24 staining and reporter expression, indicating low background noise in the absence of HIV. In all cell lines, reporter expression (GFP or mCherry) was predominantly found within the p24+ population, indicating high HIV-dependent specificity. However, there were differences in the percentage of cells responding to HIV infection, with ranges between 1.3% to 42.9% for HIV-1(NL4-3) and 0.1% to 69.9% for HIV-1(JR-CSF). These variations are likely due to clonal differences in the expression of viral receptors, reporter integration diversity, and other cellular factors. Ultimately, two of the reporter cell lines, Sup-GFP (69.9% for JR-CSF) and Rev-A3R5-GFP (42.9% for NL4-3), were identified as the most robust cells for the two lab-adapted HIV strains from this comparative study.

We tested our Rev-dependent cells for their ability to detect infection with a primary HIV-1 Clade C isolate, 27Z-P. As an example, MoltR5-GGR cells (Gaussia luciferase-GFP-Reporter) were infected with a five-fold dilution series of cell-free virus for periods ranging from 3 to 8 days (**Fig. 2A**). The percentage of infected cells was quantified through flow cytometry, while culture supernatants were assayed for GLuc activity. The resulting titration curve indicated a strong linear correlation (R^2^ = 0.9995) between the number of GFP+ cells and the dosage of the input virus (**Fig. 2B**). Similarly, the GLuc assay exhibited a straight-line correlation (**Fig. 2C**). However, we observed that undiluted culture supernatants saturated the GLuc signal at the three highest doses by day 8. For the lowest doses, the detection limits were determined to be 10L^3^ at day 3 and 10□□ at day 8. These results demonstrated that high sensitivity to detect replicating viruses can be achieved through a prolonged infection course.

**Fig. 2.**
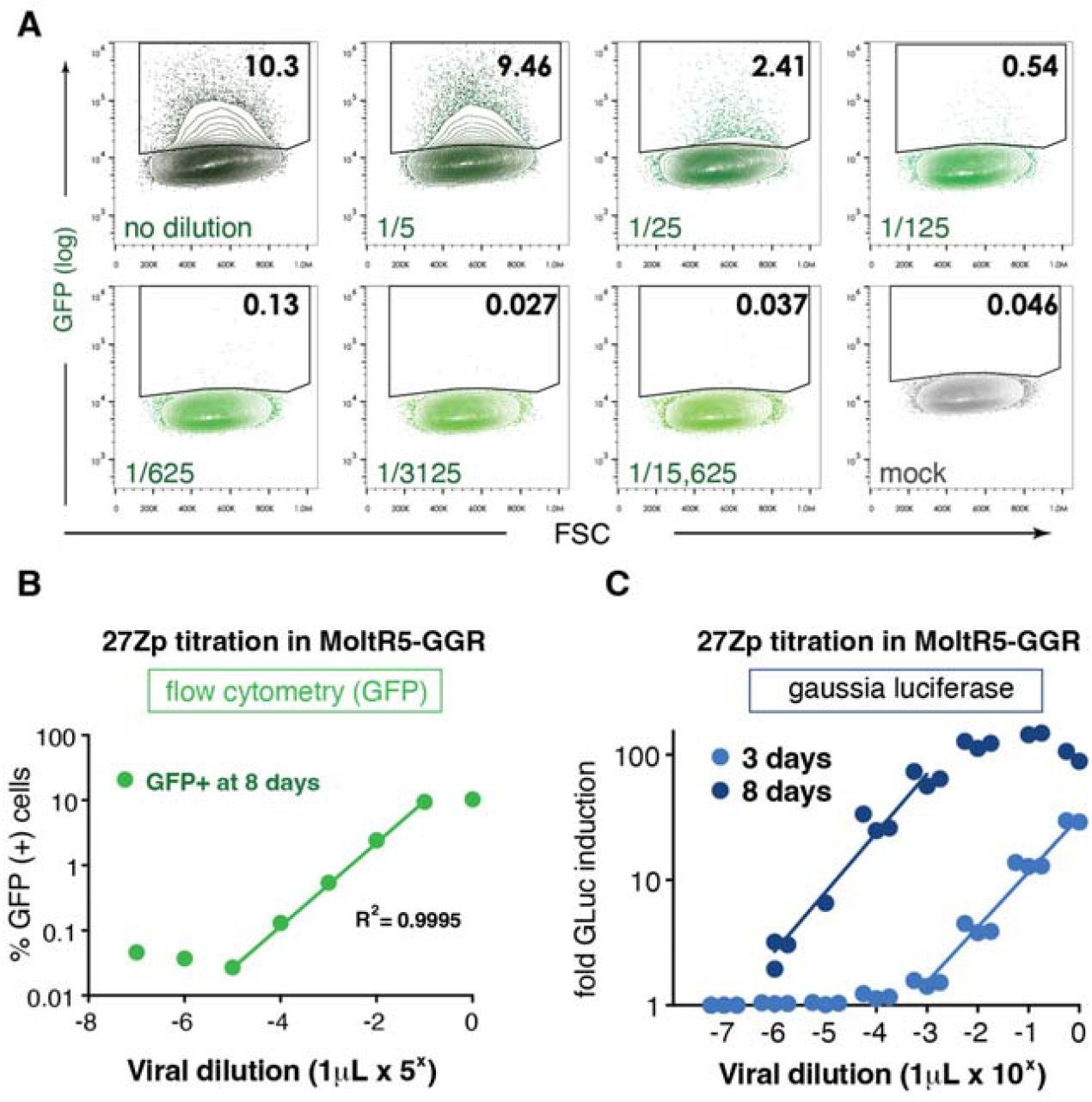
Quantitative detection of primary Clade C isolate 27Z-P in MoltR5-GG cells. (**A)** Fivefold dilution series of cell free viral stock in MoltR5-GGR cells cultured for 8 days post-infection. The GFP-positive fraction for each dilution is indicated in black (as a percentage), with the corresponding dilution indicated in green text. Before flow cytometry, cells were harvested from 96-well culture format with simple pipetting. (**B**) Plot of log percent GFP-positive fraction from (**A**) versus logs viral dilution showing high correlation in the linear region of the curve. (**C**) A similar experiment using serial tenfold dilutions of 27Z-P in MoltR5-GGR cells. Supernatants were sampled in triplicate on day 3 and day 8 to quantify GLuc activity without disturbing the cultured cells. Titration curve shows saturation at three highest doses on day 8, with absence of signal at the lowest dose.

### Validation of reporter specificity and HIV Tat- and Rev-dependency

We further validated the requirement of Tat and Rev in Rev-dependent reporter cells, using Rev-CEM as an example. Rev-CEM was infected with either wild-type HIV-1 or a Rev mutant, HIV-1(RS4X) [31]. Deleting Rev from the HIV-1 genome diminished GFP reporter expression in infected cells (**Fig. 3A**), demonstrating that other viral proteins expressed during HIV-1(RS4X) infection were insufficient to induce detectable reporter expression. In contrast, following infection with an integrase mutant HIV-1 (D116N), low-level GFP expression was detected (**Fig. 3A**). It is known that certain HIV-1 integrase mutants can mediate low-level expression of viral early proteins such as Tat, Rev, and Nef [33, 52-54]. We also constructed a lentiviral vector, pLenti-Tat/Rev-Neo, which expresses only Tat and Rev, and used the resulting lentiviral particles to infect Rev-A3R5-GFP. As shown in **Fig. 3B**, expression of Tat and Rev was sufficient to mediate GFP reporter expression in Rev-A3R5-GFP. These findings are also consistent with our prior work demonstrating that intracellular, but not extracellular Tat is sufficient to induce reporter expression in Rev-transfected TZM-gfp cells [23].

**Fig. 3.**
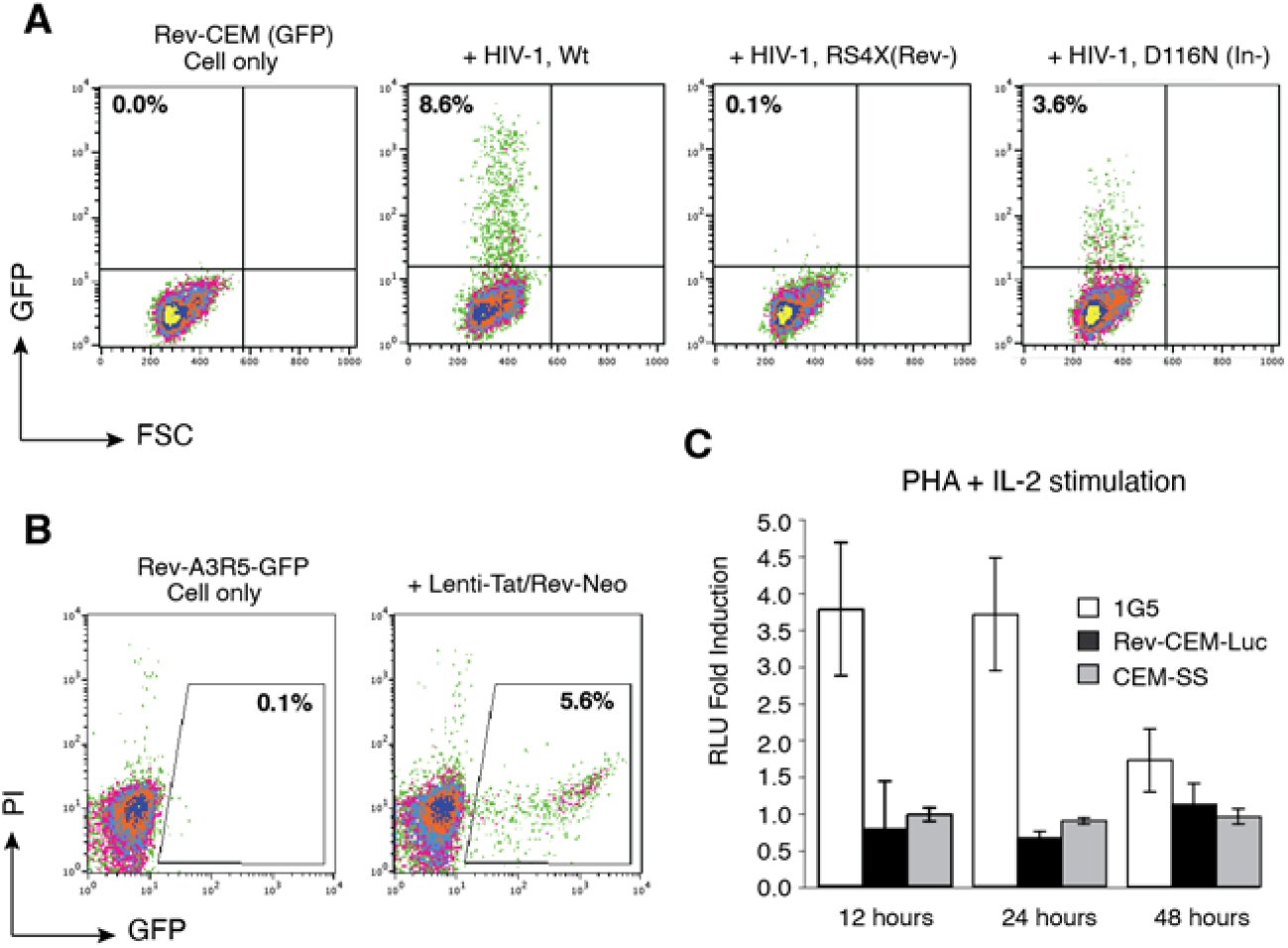
Validation of reporter specificity and HIV Tat- and Rev-dependency in Rev-dependent reporter cells. **(A)** Lack of reporter expression in the absence of a functional Rev gene. Rev-CEM cells were infected with HIV-1(NL4-3) (WT), a Rev mutant (RS4X), or an integrase mutant (D116N), using an equal level of p24 viral input. Following infection, GFP expression was quantified at 48 hours post infection. (**B**) Tat- and Rev-dependent reporter expression of reporter cells. Rev-A3R5-GFP cells were infected with a lentiviral particle, Lenti-Tat/Rev-Neo. GFP expression were quantified at 48 hours post infection. (**C**) Reporter responses to IL-2 plus PHA stimulation in 1G5 and Rev-CEM-Luc cells. Luminescence is expressed as a fold induction over untreated controls, and CEM-SS cells served as a non-luminescent control. Cells were treated with 1 U/ml IL-2 and 2 μg/ml PHA for the entirety of the indicated time points and analyzed for luciferase expression.

**Fig. 4.**
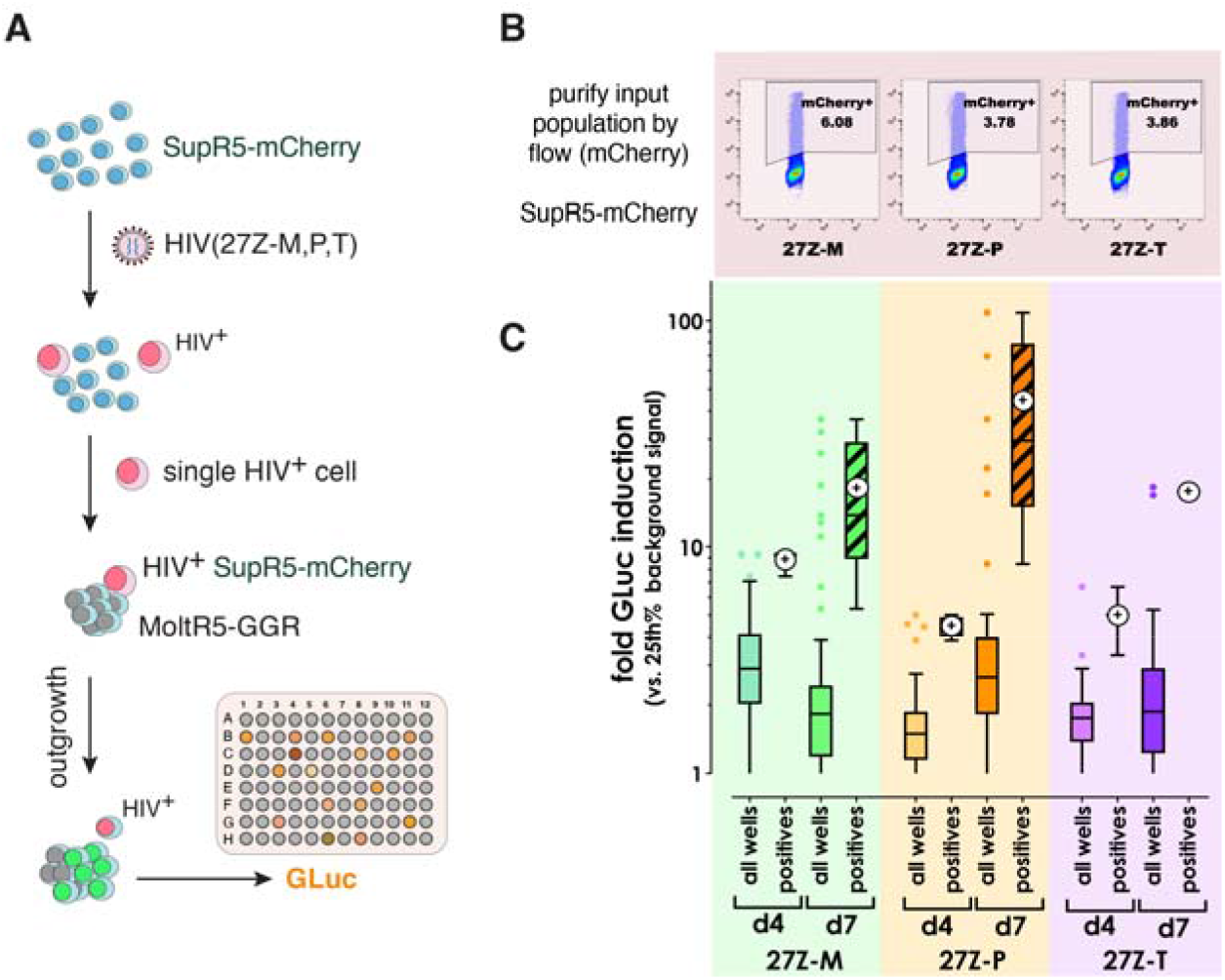
Single HIV infected cells seed a robust quantitative viral outgrowth assay in MoltR5-GG cells. (**A**) Schematic of the outgrowth assay. (**B** and **C**) SupR5-mCherry cells were infected with the indicated virus stocks and mCherry-positive cells were purified by FACS. Single mCherry-positive cells were then seeded at sub-clonal density in co-culture with MoltR5-GGR for QVOA **(C**). Tukey statistical analyses reveals outliers (> 1.5 IQR) that harbor GLuc activity. GLuc signal at days 4 and 7 of co-culture (d4, d7) for the whole seeded population is plotted for each virus and timepoint (solid color bars), with GLUc-positive outliers marked above (colored dots). Boxes represent the interquartile range (25-75 percentile), with median drawn as a straight line. Whiskers delineate the 1.5x IQ. All wells scoring positive were then re-plotted to show their distribution (hatched bars). The geometric mean is indicated by the round ‘+’ symbol.

Given the minimal requirement of Tat and Rev to turn on the reporter, we asked whether other T cell activators such as PHA plus IL-2 could activate the Rev-dependent cells non-specifically. The concomitant treatment of cells with PHA and IL-2 serves as an activating stimulus, inducing cell proliferation and the activation of transcription factors and the HIV LTR [55]. For comparison, we also tested PHA plus IL-2 stimulation of a Tat-dependent, LTR-driven luciferase reporter cell, 1G5 [34]. The Rev-dependent T cells, Rev-CEM-Luc, and 1G5 were stimulated with PHA plus IL-2 under identical conditions, and luminescence readings were collected at 12, 24, and 48 hours (**Fig. 3C**). The 1G5 cells exhibited a significant 2-4 fold increase in luciferase reporter response compared to untreated cells over the time points, while Rev-CEM-Luc showed minimal responses, similar to untreated control cells. These results demonstrate a high stringency and the requirement of both Tat and Rev for luciferase induction in Rev-dependent HIV.

### Illustration of MoltR5-GGR for highly sensitive Quantitative Viral Outgrowth Assay (QVOA) using single HIV-infected cells

The importance of HIV phenotypic diversity *in vivo* remains incompletely understood. Similarly, the determinants of *in vivo* viral fitness and cell host range remain poorly correlative with performance during viral outgrowth assays in vitro. There remains a need for a robust cellular platform to study the phenotypes of viral quasispecies and their potential impact on viral pathogenesis or persistence *in vivo*. Previously, we developed Sup-GGR cells for a lymphoid cell viral outgrowth assay [27]. Among our more recent variants of Rev-dependent reporter cells, one T cell line, MoltR5-GGR, was found to give rise to approximately 4-fold and 6-fold higher signals than Sup-GGR for R5- and X4-tropic viruses, respectively (**Fig. 1C**). This enhanced sensitivity suggests that MoltR5-GGR could offer high performance during limiting dilution quantitative viral outgrowth assay (QVOA). We further explored MoltR5-GGR in this context using HIV-positive input cells (SupR5-mCherry) harboring one of three viral swarms that were previously isolated in culture from alveolar macrophages or PBMCs of a viremic Malawian adult. These swarms contain highly related but genetically distinct viruses that were derived from separate anatomic compartments using different co-culture cellular hosts, all with the intention of capturing phenotypic diversity. Infected SupR5-mCherry cells were collected by FACS sorting, and plated at single cell density in co-culture with MoltR5-GGR. Culture supernatant was collected longitudinally at 4 and 7 days of co-culture, and replication was quantified using Gaussia luciferase assay.

Over the course of the experiment, specifically from day 4 to day 7 post co-culture, the percentage of GLuc-positive wells rose in both the 27Z-M (5.5% to 16.7%) and 27Z-P outgrowth plates (7.4% to 11.1%) (**Fig. 3B**), while for swarm 27Z-T, the assay was maximally sensitive by Day 4 (3.7% positive on both days). The GLuc assay indicated that the positive wells (outliers defined as > 1.5 IQR through B Tukey statistical analyses) exhibited an increase in mean GLuc induction between day 4 and 7 of the assay (from 8.7 to 18.2-fold for 27Z-M, from 4.5 to 43.8-fold for 27Z-P, and from 5.0 to 17.7-fold for 27Z-T (**Fig. 3C**). GLuc induction was calculated compared to the 25^th^ percentile of uninfected wells of MoltR5-GGR cells on each plate. This system affords the analysis of genetic variants in each swarm that yield strong induction through either rapid infection kinetics, modulated cytopathicity, or changes in Tat/Rev-dependent gene expression. Importantly, these results are achieved by longitudinal sampling of culture medium in the same wells over time. At the end of the assay, wells containing viral variants of interest can be harvested live for expansion, sequencing, and further phenotyping. Our findings highlight the potential of the MoltR5-GGR- and other Rev-dependent reporter based QVOA assays to detect and characterize viruses from single infected cells early during outgrowth.

### Illustration of Rev-A3R5-GFP and Rev-CEM-Luc for screening and quantifying neutralizing antibodies

The A3R5 lymphoblastic T cell was developed by McLinden *et al.* [56], based on the human CD4^+^/CXCR4^+^/a4b7^+^ T-lymphoblastoid cell line A3.01[29]. A3R5 cells express CD4, CXCR4, and CCR5, with notably lower CCR5 surface expression compared to TZM-bl but similar levels as stimulated PBMC [56]. This cell line is susceptible to infection with a variety of CXCR4- and CCR5-tropic circulating strains of HIV-1. Notably, neutralization mediated by a diverse panel of monoclonal antibodies and HIV^+^ sera has shown consistently greater efficacy in A3R5 compared to TZM-bl cells [56]. A3R5 is particularly useful for detecting weak neutralizing antibody responses against Tier 2 strains of HIV-1 and has undergone extensive optimization and validation under Good Clinical Laboratory Practice (GCLP) conditions. The validated A3R5 assay has been employed to assess vaccine-elicited neutralizing antibodies and to identify correlates of protection during vaccine trials worldwide [57]. However, the assay requires Env gene cloning and pseudotyping a replication-competent reporter virus, when testing neutralization against specific new viral mutants or strains. We took advantage of these unique features of A3R5, and embedded the Rev-dependent GFP reporter (pNL-GFP-RRE-SA-puro) into A3R5 to establish Rev-A3R5-GFP reporter cells. This cell line is particularly suited to direct quantification of antibody neutralization against replication competent HIV-1, either laboratory-adapted or primary stains. As a proof-of-concept, we performed a neutralizing antibody screening using HIV-1(NL4-3) infection of Rev-A3R5-GFP (**Fig. 5A**). Prior to infection, HIV-1 viral stocks were incubated with or without individual Anti-Env antibodies from a screening panel. After 1 hour of neutralization, the antibody-HIV-1 mixture was used to infect Rev-A3R5-GFP cells for two hours, followed by washing, and culturing for two days prior to flow cytometric analysis. As summarized in **Fig. 5A,** the results indicated that the strongest neutralizing antibodies were IgG1 b12, VRC03, 5F7, 17b, 654 30D, F425 A1g8, and HJ16, while the remaining antibodies demonstrated weak to non-neutralizing activity. The IC50 value for VRC03 was quantified at 1 µg/ml (**Fig. 5B**).

**Fig. 5.**
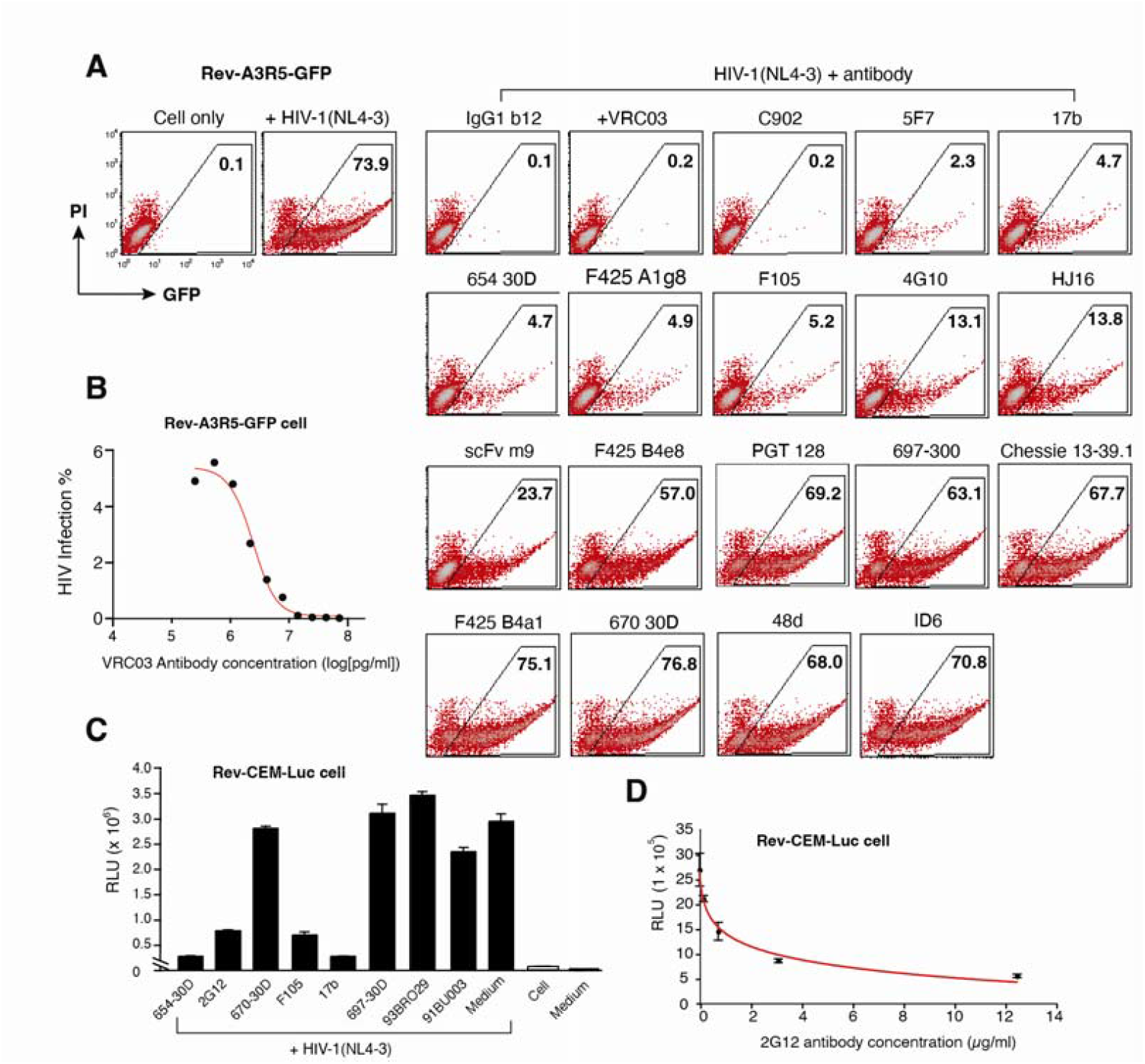
Screening and Quantification of anti-HIV antibodies with Rev-dependent reporter cells. (A) Screening for the neutralization activity of anti-HIV antibodies using Rev-A3R5-GFP. HIV-1(NL4-3) were incubated with or without anti-HIV antibodies for 1 hour, and then used to infect Rev-A3R5-GFP cells for 2 hours. Cells were washed, cultured for 48 hours, and then analyzed for GFP expression. (**B**) Quantification of the neutralization activity of the anti-HIV monoclonal antibody VRC03 using Rev-A3R5-GFP. Serial dilutions of VRC03 were incubated with HIV-1(NL4-3) for 2 hours, added to Rev-A3R5-GFP cells, and incubated for 48 hours. Cells were washed, cultured for 48 hours, and then analyzed for GFP expression. The antibody dose-dependent neutralization curve is plotted. (**C**) Screening for the neutralization activity of anti-HIV antibodies using Rev-CEM-Luc. Five micrograms of indicated antibodies were incubated with HIV-1(NL4-3) for 2 hours, and then added to Rev-CEM-Luc and incubated for 48 h. The polyclonal, serum-derived antibodies, 93BRO29 and 91BU003, were diluted 1:50 in medium. Uninfected cells or medium alone were also included as controls. Inhibition of viral infection is indicated by a reduction in relative luciferase activity (RLU) as compared to mock-treated controls. **(D)** Quantification of the neutralization activity of the anti-HIV monoclonal antibody 2G12 using Rev-CEM-Luc. Serial four-fold dilutions of 50 μg/ml stock solution were incubated with HIV-1(NL4-3) for 2 hours, added to Rev-CEM-Luc cells, and incubated for 48 hours for reporter luminescence. The red line is the best-fit curve of exponential decay, with a calculated IC_50_ of 0.94 µg/ml.

We repeated this screen in another Rev-dependent cell line, Rev-CEM-Luc, with selected antibodies neutralizing HIV-1(NL4-3). Among the tested antibodies, 654-30D, 2G12, F105, and 17b exhibited strong inhibition of viral infection (**Fig. 5C**). The antibody 2G12 had an IC_50_ against HIV-1(NL4-3) measured at 0.98±0.42 µg/ml in a previous study [58], compared to an IC_50_ calculated at 0.94 µg/ml (**Fig. 5D**) in our assay. This illustrates the ability of Rev-CEM-Luc cells to reproducibly quantify neutralizing antibodies consistent with existing literature using full length, replication competent virus.

### Illustration of Rev-A3R5-GFP for quantifying the potency of HIV restriction factors

HIV restriction factors such as PSGL-1 [50, 59, 60], APOBEC3G (A3G) [61], and SERINC5 [62, 63] diminish infectivity when incorporated into virions. The inhibition of virion infectivity is frequently dose-dependent and can be quantified using indicator cells. We used Rev-A3R5-GFP to accurately gauge the efficacy of these restriction factors. One significant benefit of employing Rev-dependent cells is that reporter expression or inhibition is intricately regulated based on the ability of infecting virions to express Tat and Rev in target reporter cells. In contrast, non-infectious particles and cell-free viral proteins like extracellular Tat or Env found in virus preparations cannot activate the reporter [23]. This provides a key advantage over LTR-driven reporter cells and allows for ultra-sensitive detection of virion inactivation at low concentration of restriction factors. Using PSGL-1 as an example, Rev-A3R5-GFP enabled the detection of virion restriction at concentrations as low as 0.5-1 ng of PSGL-1 (**Fig. 6A**) [50]. The dosage-dependent inhibition of PSGL-1 allowed us to determine its IC_50_ to be 3.64 ng in this particular set of quantification (**Fig. 6B**). We also compared the anti-HIV-1 activities of APOBEC3G and SERINC5, determining that SERINC5 exhibited a low IC_50_ of 4.63 ng, a similar potency to PSGL-1 against wild-type HIV, while APOBEC3G exhibited a much higher IC_50_ of 380.52 ng against HIV-1(NL4-3) (**Fig. 6B**). These findings agree with prior work showing that APOBEC3G is more effective against Vif-negative viruses but less so against wild-type HIV-1 when Vif is present [61].

**Fig. 6.**
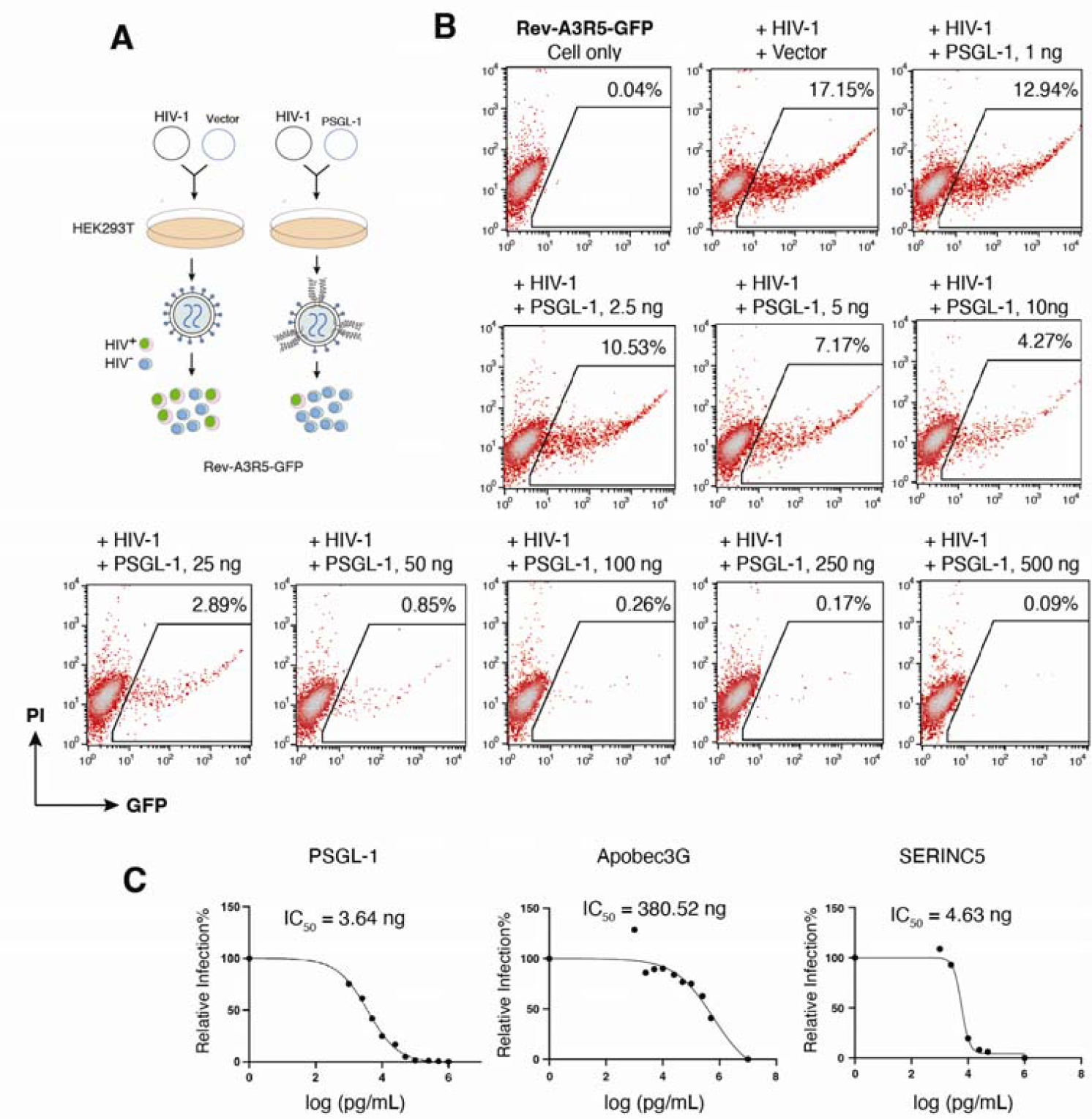
Accurate quantifying of anti-HIV restriction factors using Rev-A3R5-GFP. **(A)** Schematic of the assay for quantifying the antiviral activity of an anti-HIV restriction factor PSGL-1 using Rev-A3R5-GFP. (**B**) PSGL-1 dose-dependent inhibition of HIV infectivity quantified using Rev-A3R5-GFP. HEK293T cells were cotransfected with 1 μg pNL4-3 DNA plus various amounts of PSGL-1 DNA (1 to 500 ng). Virions were collected at 48 hours post-cotransfection and used to infect Rev-A3R5-GFP, using an equal p24 level of virus input. Virus infectivity was quantified with flow cytometry of HIV+ cells (GFP+) at 48 hours post infection. The 50% inhibitory dosage (IC_50_) was determined and plotted in (**C**). (**C**) Restriction factors such as APOBEC3G and SERINC5 were similarly quantified for their IC_50_ using Rev-A3R5-GFP.

## Discussion

In this work, we exhibited the utility of several Rev-dependent reporter cell lines. The first such reporter systems were developed by Wu et al. [17, 18]; recently, Gludish and Salasc et al. expanded the system by developing new reporter cells in several CXCR4 and CCR5 dual-expressing lines [23, 27]. We also provide the first characterization of Rev-dependent reporter cells derived from MoltR5, CEM-SS, and A3R5. In these systems and superimposed on the partial intracellular Tat-dependent transcription from the LTR, an additional layer of post-transcriptional regulation is mediated by intracellular Rev binding to the RRE [21, 64] enabling nuclear export of unspliced, reporter ORF-containing transcripts. By imposing a Boolean AND requirement for intracellular Tat and Rev, reporter expression is rendered nonresponsive to extraneous sources of stimulation, including mitogens: this was shown in Rev-CEM-Luc, where addition of PHA and IL-2 induced reporter expression in the LTR-driven luciferase reporter line and Jurkat derivative, 1G5 [34], but not Rev-CEM-Luc. Presumably, in situations where Tat-independent transcription is driven by high levels of endogenous transcriptional transactivators, like NF-κB [65], NFAT [66], SP-1 [67], and AP-1 [68, 69], Rev-dependent reporter systems produce full-length transcripts containing the reporter ORF, but these are sequestered in the nucleus by established mechanisms [70, 71], preventing expression. The requirement for intracellular Rev appears to be nearly absolute in physiological contexts; however, Gludish et al. [23]exhibited that supraphysiological levels of intracellular Tat can induce low levels of reporter expression. In these situations, it is plausible that nuclear retention mechanisms can become overwhelmed, resulting in leakage of unspliced transcripts, which are sufficiently stable and persistent in the cytoplasm to enable translation. Interestingly, this effect was not observed with extracellular Tat, implying that inefficiencies in cellular entry, nuclear localization, or the acquisition of specific post-translational modifications might preclude sufficient transactivation [23]. However, we also demonstrated that, under normal infection conditions, Rev-independent reporter expression is unlikely, as a ΔRev mutant derived from HIV-1 NL4-3 failed to induce reporter expression [31].

Additionally, we established the permissiveness and responsiveness of Rev-dependent cell lines to the X4 and R5-tropic viruses, HIV-1NL4-3 and HIV-1JR-CSF. While all lines demonstrated HIV-driven reporter gene expression, considerable variability was observed. This is likely attributable to clone-specific factors that were not explored in this work. Probable drivers mediating variable responsiveness are coreceptor and restriction factor expression, while integration site location and copy number presumably contribute to sensitivity, and the combination of these and other factors impact maximal reporter expression in the presence of higher viral titers. The low-level responsiveness to R5-tropic HIV-1 in Rev-CEM-Luc and Rev-CEM suggests minimal CCR5 expression: while CEM cells typically express undetectable-to-negligible levels of CCR5, induction can occur in the presence of KLF2 [72], and it is plausible a similar effect drives de minimis expression in these clones. Among the reporter lines assayed, SupR5-GFP and Rev-A3R5-GFP exhibited the highest responses to HIV-1JR-CSF and HIV-1NL4-3, respectively, indicating a singular combination of the above-mentioned factors contributed to heightened responsiveness.

The sensitivity of Rev-dependent reporter systems was specifically addressed in MoltR5-GGR cells, which express hrGFP and secreted gaussia luciferase, using a primary HIV-1 clade C isolate, 27Z-P, derived from an uncharacterized viral swarm and developed by Gludish et al. [23]. High-level sensitivity to cell-free virus was demonstrated for both reporters by viral dilution assay, with luciferase endpoints being more sensitive. Moreover, the facile longitudinal sampling capabilities afforded by secreted luciferase allowed for more sensitive detection at later timepoints. However, the MoltR5-GGR system was designed to measure infection in co-culture systems in VOAs. As such, to determine functionality in this application where a single cell mediates infection, MoltR5-GGR cells were exposed to a single infected SupR5-mCherry per well. The development of reporter-positive wells (both GFP fluorescence and GLuc luminescence) enables flexible interrogation of phenotype and associated viral genotypes for individual clones among swarms isolated from human patients. Here, we compared strains from outgrowth culture derived from bronchoalveolar lavage and the peripheral blood of a viremic adult patient from Blantyre, Malawi. This unique use case for MoltR5-GGR affords the analysis of primary isolates prior to laboratory adaptation, enabling the identification of *in situ* HIV genotypes, including unique env sequences that may be otherwise lost during culture: the characterization of these genotypes could establish novel evolutionary tradeoffs, host factor dependencies, and interactions that may be superfluous *in vitro*.

Lastly, we explored the functionality of the Rev-dependent systems in another salient application, the identification and screening of neutralizing antibodies and host restriction factors. Using Rev-A3R5-GFP and Rev-CEM-Luc, dose-dependent neutralization was observed with antibodies, VRC03 and 2G12, respectively: in both cases, the observed IC_50_ was consistent with reported values from Huskens et al. [58] and a commercial VRC03 vendor. Lastly, suppression of viral replication by host restriction factors in Rev-A3R5-GFP was demonstrated, where PSGL-1 [59], APOBEC3G [61], and SERINC5 [62, 63] inhibited reporter expression dose-dependently after transfection into producer cells. The sensitive detection of infection inhibition by low levels of virion-incorporated restriction factors and by neutralizing antibodies highlights the utility of these systems in screening and characterization. Furthermore, while not explored in this work, we have previously demonstrated small molecule antiviral screening using Rev-CEM-Luc [73].

Cumulatively, the data presented herein establishes the sensitivity, specificity, and facility of Rev-dependent reporter cell systems, while also introducing three new Rev-dependent reporter lines. Collectively, these reporter lines constitute a versatile toolkit for HIV research. In requiring both intracellular Tat and Rev, these systems are uniquely capable of assessing viral replication and its inhibition in complex matrices that may contain mitogens or other transcription modulating components. Additionally, these systems allow for the interrogation of novel genotypes that may exist only transiently in primary isolates. Furthermore, these systems are highly functional in the screening and characterization of neutralizing antibodies, restriction factors, and antivirals. Given this, the development and dissemination of these reporter cell lines is broadly substantial to HIV research.

## Materials and Methods

### DNA Vectors

HIV-1 DNA vectors, pHIV-1(NL4-3), pHIV-1(RS4X) mutant, and pHIV-1(NL4-3_D116N_) were obtained from the NIH AIDS Reagent Program. The lentiviral vectors pNL-RRE-SA, pNL-GFP-RRE, pNL-Luc-RRE have been described previously [17, 24, 32]. pNL-GFP-RRE-Puro was produced by Gibson Assembly of the puromycin resistance gene into pNL-GFP-RRE. pNL-GGR-RRE (SA) was derived by cloning a GLuc cassette (synthesized by IDT Inc.) on a SmaI-SphI fragment into pNL-GFP-RRE(SA) cut with the same enzymes. Construction of pHIV-nef-IRES-mCherry has been described previously [23]. pcDNA3.1-PSGL-1 and pcDNA3.1-A3G were synthesized by Twist Bioscience, using the pcDNA3.1 vector. pCMV6-SERINC5 vector was kindly provided by Dr. Yonghui Zheng. pLenti-Tat/Rev-Neo was synthesized based on the HIV-1(NL4-3) sequence. All HIV-1 ORFs except Tat and Rev are deleted, and the vector contains splicing donor and receptor sites (SD1 to SA7) and RRE. The *nef* gene region is replaced with the neo gene. The vector express Tat and Rev from the HIV-1 LTR promoter, and Neo from the hPKG promoter.

### Cell Lines and Viruses

HEK293T (ATCC) cells were maintained in Dulbecco’s modified Eagle’s medium (DMEM) (Invitrogen) containing 10% heat-inactivated FBS and 100 units/ml of penicillin and 100 µg/ml of streptomycin (Invitrogen). T cell line 1G5 [34](NIH AIDS Reagent Program) and the Rev-dependent indicator T cells, Rev-CEM, Rev-CEM-Luc, Rev-A3-GFP/Luc, Rev-A3R5-GFP, Rev-A3R5-GFP/Luc, and LTR-luc, were cultured in RPMI-1640 plus 10% FBS supplemented with 100 units/ml of penicillin and 100 µg/ml of streptomycin (Invitrogen). Rev-CEM-Luc and 1G5 were treated with PHA and IL-2 in a 24-well plate, one million cells were treated with PHA and IL-2 (R&D Systems) simultaneously at a final concentration of 2 μg/ml and 1 U/ml, respectively. For HIV-1 virus production, HEK293T cells were transfected with 1 μg of HIV-1 DNA (pNL4-3) or cotransfected with 1 μg of HIV-1 DNA (pNL4-3) plus the indicated doses of pcDNA3.1-PSGL-1, pcDNA3.1-A3G, or pCMV6-SERINC5, in 6-well plates using transfection reagent EZ-Fectin (Virongy) as recommended by the manufacturer. Virion-containing supernatants were collected at 48 hours post transfection. SupR5 and M4R5 cells were each transduced with the GGR lentiviral vector (Fig 1A) and a panel of clones were generated by serial dilution. Individual clones were replica plated and selected for robust reporter gene expression by flow cytometry after infection with HIV-nef-IRES-mCherry/BaL env.

### Reporter Quantification by Flow Cytometry and Luciferase Assay

GFP cells were analyzed using a FACSCalibur flow cytometer (BD Biosciences) and CellQuest software (BD Biosciences). Luciferase assay was conducted using a commercial luciferase assay kit (Virongy) using procedures as recommended by the manufacturer. Briefly, cells were pelleted at 1,200 rpm for 5 minutes and lysed in 300 μl lysis buffer (Virongy), and luciferin substrate (Virongy) was injected into each well of reading plates utilizing a GloMax-Multi luminometer with the Single Injector System (Promega).

### Virus Neutralization

HIV positive patient plasma 91BU003 (infected with a clade C virus), 93BR029 (infected with a clade B virus), and monoclonal antibodies including anti-HIV-1 gp120 monoclonal (VRC03), anti-HIV-1 IIIB gp160 monoclonal antibody (Chessie 13-39.1), anti-HIV-1 gp120 monoclonal (48d), anti-HIV gp120 monoclonal (654-30D), HIV-1 gp120 monoclonal Antibody (F425 B4e8), anti-HIV-1 gp120 monoclonal (4G10), anti-HIV-1 gp120 monoclonal (5F7), anti-HIV-1 gp120 monoclonal (670-30D), HIV-1 gp120 monoclonal antibody (F425 B4a1), anti-HIV-1 gp120 monoclonal (scFv m9), anti-HIV-1 gp120 monoclonal (IgG1 b12), HIV-1 gp120 monoclonal Antibody (ID6), anti-HIV-1 gp120 monoclonal (PGT128), anti-HIV-1 gp120 monoclonal (C902), anti-HIV-1 gp120 monoclonal (F105), anti-HIV-1-gp120 monoclonal (F425 A1g8), anti-HIV-1 gp120 monoclonal (HJ16), anti-HIV-1 gp120 monoclonal (2G12), anti-HIV-1 gp120 monoclonal (697-30D), anti-HIV-1 gp120 monoclonal (17b), 93BR029, and 91BU003 were received from NIH AIDS Reagent Program. Individual antibodies were incubated with HIV-1(NL4-3) virus for 1 hour at 37 ºC prior to addition to one million Rev-A3R5-GFP or Rev-CEM-Luc cells. Cells were infected for 2 hours, washed, and then analyzed by flow cytometry or luciferase assay at 48 hours post-infection.

### Single cell quantitative viral outgrowth assay

Using three separate viral supernatants derived from a Malawian adult (27Z), SupR5-mCherry cells were spinoculated at 2200 rpm for 90 minutes at MOI of 0.05 in the presence of DEAE-dextran. Cells were washed three times the following day and replated in fresh medium. At 48 hours post-infection, cells were harvested and purified by FACS on a BioRad S3e instrument. Collected cells were counted and diluted for seeding at sub-clonal density (∼0.3 cells/well), then placed in co-culture with previously seeded MoltR5-GGR cells in U-bottom 96-well culture dishes, total volume 200uL. Outgrowth infection was monitored by fluorescence microscopy (Leica SP5). Medium was harvested without mixing at 4 days and 7 days of co-culture, and stored at -80°C. Samples were thawed for Gaussia assay (Gaussia Glow, Pierce Cat. No. 16161) according to the manufacturer instructions and acquired on an EnVision instrument (Perkin Elmer/Revvity). Background subtracted raw data were analyzed in Microsoft Excel and Graphpad Prism software for Tukey outliers and box-whisker plots comparing each well for GLuc induction versus the 25^th^ percentile GLuc induction in negative control wells (no infected input cells).

## Acknowledgments

The authors wish to thank the NIH HIV Reagent Program for reagents. This work was supported by NIH awards K01OD031968 and R21OD037879 (to D.W.G), R01AI148012 (to Y.W.), AI176575 (to D.G.R), and Wellcome Trust Clinical Intermediate Fellowship 088696/Z/09/Z (to H.C.M.).

## Author contributions

D.W.G and Y.W. conceived and supervised the study. D.W.G, Y.W. performed experimental data analysis. D.G.R. and H.C.M. contributed clinical isolates. M.S., J.C., B.H, T.T.V., J.H.L, G.L.B, Y.H., H.L., J.G., D.Y., S.I. performed experiments. M.S., Y.W., and D.W.G. wrote the original manuscript draft. All authors reviewed the manuscript.

## References

1. Felber BK, Pavlakis GN. A quantitative bioassay for HIV-1 based on trans-activation. Science. 1988;239(4836):184–7. doi: 10.1126/science.3422113. PubMed PMID: 3422113.

2. Schwartz S, Felber BK, Fenyö EM, Pavlakis GN. Rapidly and slowly replicating human immunodeficiency virus type 1 isolates can be distinguished according to target-cell tropism in T-cell and monocyte cell lines. Proc Natl Acad Sci U S A. 1989;86(18):7200–3. doi: 10.1073/pnas.86.18.7200. PubMed PMID: 2789383; PubMed Central PMCID: PMCPMC298024.

3. Rocancourt D, Bonnerot C, Jouin H, Emerman M, Nicolas JF. Activation of a beta-galactosidase recombinant provirus: application to titration of human immunodeficiency virus (HIV) and HIV-infected cells. J Virol. 1990;64(6):2660–8. doi: 10.1128/jvi.64.6.2660-2668.1990. PubMed PMID: 2110596; PubMed Central PMCID: PMCPMC249444.

4. Kimpton J, Emerman M. Detection of replication-competent and pseudotyped human immunodeficiency virus with a sensitive cell line on the basis of activation of an integrated beta-galactosidase gene. J Virol. 1992;66(4):2232-9. PubMed PMID: 1548759.

5. Platt EJ, Wehrly K, Kuhmann SE, Chesebro B, Kabat D. Effects of CCR5 and CD4 cell surface concentrations on infections by macrophagetropic isolates of human immunodeficiency virus type 1. J Virol. 1998;72(4):2855-64. PubMed PMID: 9525605.

6. Muesing MA, Smith DH, Capon DJ. Regulation of mRNA accumulation by a human immunodeficiency virus trans-activator protein. Cell. 1987;48(4):691–701. doi: 10.1016/0092-8674(87)90247-9. PubMed PMID: 3643816.

7. Dingwall C, Ernberg I, Gait MJ, Green SM, Heaphy S, Karn J, et al. Human immunodeficiency virus 1 tat protein binds trans-activation-responsive region (TAR) RNA in vitro. Proc Natl Acad Sci U S A. 1989;86(18):6925–9. doi: 10.1073/pnas.86.18.6925. PubMed PMID: 2476805; PubMed Central PMCID: PMCPMC297963.

8. Berkhout B, Silverman RH, Jeang KT. Tat trans-activates the human immunodeficiency virus through a nascent RNA target. Cell. 1989;59(2):273–82.

9. Kao SY, Calman AF, Luciw PA, Peterlin BM. Anti-termination of transcription within the long terminal repeat of HIV-1 by tat gene product. Nature. 1987;330(6147):489–93.

10. Laspia MF, Rice AP, Mathews MB. HIV-1 Tat protein increases transcriptional initiation and stabilizes elongation. Cell. 1989;59(2):283–92. doi: 10.1016/0092-8674(89)90290-0. PubMed PMID: 2553266.

11. Kato H, Sumimoto H, Pognonec P, Chen CH, Rosen CA, Roeder RG. HIV-1 Tat acts as a processivity factor in vitro in conjunction with cellular elongation factors. Genes Dev. 1992;6(4):655–66. doi: 10.1101/gad.6.4.655. PubMed PMID: 1559613.

12. Raha T, Cheng SW, Green MR. HIV-1 Tat stimulates transcription complex assembly through recruitment of TBP in the absence of TAFs. PLoS Biol. 2005;3(2):e44. Epub 20050208. doi: 10.1371/journal.pbio.0030044. PubMed PMID: 15719058; PubMed Central PMCID: PMCPMC546330.

13. Brady J, Kashanchi F. Tat gets the “green” light on transcription initiation. Retrovirology. 2005;2:69. Epub 20051109. doi: 10.1186/1742-4690-2-69. PubMed PMID: 16280076; PubMed Central PMCID: PMCPMC1308864.

14. Gervaix A, West D, Leoni LM, Richman DD, Wong-Staal F, Corbeil J. A new reporter cell line to monitor HIV infection and drug susceptibility in vitro. Proc Natl Acad Sci U S A. 1997;94(9):4653–8.

15. Siekevitz M, Josephs SF, Dukovich M, Peffer N, Wong-Staal F, Greene WC. Activation of the HIV-1 LTR by T cell mitogens and the trans-activator protein of HTLV-I. Science. 1987;238(4833):1575–8.

16. Swingler S, Easton A, Morris A. Cytokine augmentation of HIV-1 LTR-driven gene expression in neural cells. AIDS Res Hum Retroviruses. 1992;8(4):487-93. PubMed PMID: 1599755.

17. Wu Y, Beddall MH, Marsh JW. Rev-dependent lentiviral expression vector. Retrovirology. 2007;4(1):12. PubMed PMID: 17286866.

18. Wu Y, Beddall MH, Marsh JW. Rev-dependent indicator T cell line. Current HIV Research. 2007;5:395–403.

19. Feinberg MB, Jarrett RF, Aldovini A, Gallo RC, Wong-Staal F. HTLV-III expression and production involve complex regulation at the levels of splicing and translation of viral RNA. Cell. 1986;46(6):807–17. doi: 10.1016/0092-8674(86)90062-0. PubMed PMID: 3638988.

20. Malim MH, Hauber J, Fenrick R, Cullen BR. Immunodeficiency virus rev trans-activator modulates the expression of the viral regulatory genes. Nature. 1988;335(6186):181–3. doi: 10.1038/335181a0. PubMed PMID: 3412474.

21. Malim MH, Hauber J, Le SY, Maizel JV, Cullen BR. The HIV-1 rev trans-activator acts through a structured target sequence to activate nuclear export of unspliced viral mRNA. Nature. 1989;338(6212):254–7.

22. Emerman M, Vazeux R, Peden K. The rev gene product of the human immunodeficiency virus affects envelope-specific RNA localization. Cell. 1989;57(7):1155–65. doi: 10.1016/0092-8674(89)90053-6. PubMed PMID: 2736624.

23. Gludish DW, Boliar S, Caldwell S, Tembo DL, Chimbayo ET, Jambo KC, et al. TZM-gfp cells: a tractable fluorescent tool for analysis of rare and early HIV-1 infection. Scientific Reports. 2020;10(1):19900. doi: 10.1038/s41598-020-76422-6.

24. Wang Z, Tang Z, Zheng Y, Yu D, Spear M, Iyer SR, et al. Development of a nonintegrating Rev-dependent lentiviral vector carrying diphtheria toxin A chain and human TRAF6 to target HIV reservoirs. Gene Ther. 2010;17(9):1063-76. PubMed PMID: 20410930.

25. Hetrick B, Siddiqui S, Spear M, Guo J, Liang H, Fu Y, et al. Suppression of viral rebound by a Rev-dependent lentiviral particle in SIV-infected rhesus macaques. Gene Therapy. 2024. doi: 10.1038/s41434-024-00467-9.

26. Chikere K, Webb NE, Chou T, Borm K, Sterjovski J, Gorry PR, et al. Distinct HIV-1 entry phenotypes are associated with transmission, subtype specificity, and resistance to broadly neutralizing antibodies. Retrovirology. 2014;11(1):48. doi: 10.1186/1742-4690-11-48.

27. Salasc F, Gludish DW, Jarvis I, Boliar S, Wills MR, Russell DG, et al. A novel, sensitive dual-indicator cell line for detection and quantification of inducible, replication-competent latent HIV-1 from reservoir cells. Scientific Reports. 2019;9(1):19325. doi: 10.1038/s41598-019-55596-8.

28. Means RE, Matthews T, Hoxie JA, Malim MH, Kodama T, Desrosiers RC. Ability of the V3 loop of simian immunodeficiency virus to serve as a target for antibody-mediated neutralization: correlation of neutralization sensitivity, growth in macrophages, and decreased dependence on CD4. J Virol. 2001;75(8):3903–15. doi: 10.1128/jvi.75.8.3903-3915.2001. PubMed PMID: 11264379; PubMed Central PMCID: PMCPMC114881.

29. Folks T, Benn S, Rabson A, Theodore T, Hoggan MD, Martin M, et al. Characterization of a continuous T-cell line susceptible to the cytopathic effects of the acquired immunodeficiency syndrome (AIDS)-associated retrovirus. Proc Natl Acad Sci U S A. 1985;82(13):4539-43. PubMed PMID: 2989831; PubMed Central PMCID: PMCPMC391138.

30. Nara PL, Hatch WC, Dunlop NM, Robey WG, Arthur LO, Gonda MA, et al. Simple, rapid, quantitative, syncytium-forming microassay for the detection of human immunodeficiency virus neutralizing antibody. AIDS Res Hum Retroviruses. 1987;3(3):283-302. PubMed PMID: 3481271.

31. Riggs NL, Guatelli JC. Production and characterization of high-titer stocks of rev-defective HIV-1. Virology. 1996;217(2):602-6. PubMed PMID: 8610453.

32. Young J, Tang Z, Yu Q, Yu D, Wu Y. Selective killing of HIV-1-positive macrophages and T Cells by the Rev-dependent lentivirus carrying anthrolysin O from Bacillus anthracis. Retrovirology. 2008;5(1):36. PubMed PMID: 18439272.

33. Iyer SR, Yu D, Biancotto A, Margolis LB, Wu Y. Measurement of human immunodeficiency virus type 1 preintegration transcription by using Rev-dependent Rev-CEM cells reveals a sizable transcribing DNA population comparable to that from proviral templates. J Virol. 2009;83(17):8662-73. PubMed PMID: 19553325.

34. Aguilar-Cordova E, Chinen J, Donehower L, Lewis DE, Belmont JW. A sensitive reporter cell line for HIV-1 tat activity, HIV-1 inhibitors, and T cell activation effects. AIDS Res Hum Retroviruses. 1994;10(3):295-301. PubMed PMID: 8018390.

35. Garcia JA, Wu FK, Mitsuyasu R, Gaynor RB. Interactions of cellular proteins involved in the transcriptional regulation of the human immunodeficiency virus. EMBO Journal. 1987;6(12):3761–70.

36. Swingler S, Morris A, Easton A. Tumour necrosis factor alpha and interleukin-1 beta induce specific subunits of NFKB to bind the HIV-1 enhancer: characterisation of transcription factors controlling human immunodeficiency virus type 1 gene expression in neural cells. Biochem Biophys Res Commun. 1994;203(1):623-30. PubMed PMID: 8074713.

37. Akan E, Chang-Liu CM, Watanabe J, Ishizawa K, Woloschak GE. The effects of vinblastine on the expression of human immunodeficiency virus type 1 long terminal repeat. Leuk Res. 1997;21(5):459-64. PubMed PMID: 9225075.

38. Sweet MJ, Hume DA. RAW264 macrophages stably transfected with an HIV-1 LTR reporter gene provide a sensitive bioassay for analysis of signalling pathways in macrophages stimulated with lipopolysaccharide, TNF-alpha or taxol. J Inflamm. 1995;45(2):126-35. PubMed PMID: 7583358.

39. Kurata S-i. Sensitization of the HIV-1-LTR upon Long Term Low Dose Oxidative Stress*. Journal of Biological Chemistry. 1996;271(36):21798–802. doi: 10.1074/jbc.271.36.21798.

40. Iordanskiy S, Van Duyne R, Sampey GC, Woodson CM, Fry K, Saifuddin M, et al. Therapeutic doses of irradiation activate viral transcription and induce apoptosis in HIV-1 infected cells. Virology. 2015;485:1–15. doi: 10.1016/j.virol.2015.06.021. PubMed PMID: 26184775; PubMed Central PMCID: PMCPMC4619158.

41. Merzouki A, Patel P, Cassol S, Ennaji M, Tailor P, Turcotte FR, et al. HIV-1 gp120/160 expressing cells upregulate HIV-1 LTR directed gene expression in a cell line transfected with HIV-1 LTR-reporter gene constructs. Cell Mol Biol (Noisy-le-grand). 1995;41(3):445-52. PubMed PMID: 7580840.

42. Lu HK, Gray LR, Wightman F, Ellenberg P, Khoury G, Cheng W-J, et al. Ex Vivo Response to Histone Deacetylase (HDAC) Inhibitors of the HIV Long Terminal Repeat (LTR) Derived from HIV-Infected Patients on Antiretroviral Therapy. PLOS ONE. 2014;9(11):e113341. doi: 10.1371/journal.pone.0113341.

43. Sloan RD, Kuhl BD, Donahue DA, Roland A, Bar-Magen T, Wainberg MA. Transcription of preintegrated HIV-1 cDNA modulates cell surface expression of major histocompatibility complex class I via Nef. J Virol. 2011;85(6):2828-36. PubMed PMID: 21209113.

44. Sloan RD, Donahue DA, Kuhl BD, Bar-Magen T, Wainberg MA. Expression of Nef from unintegrated HIV-1 DNA downregulates cell surface CXCR4 and CCR5 on T-lymphocytes. Retrovirology. 2011;7:44. PubMed PMID: 20465832.

45. Shuck-Lee D, Chang H, Sloan EA, Hammarskjold ML, Rekosh D. Single-nucleotide changes in the HIV Rev-response element mediate resistance to compounds that inhibit Rev function. J Virol. 2011;85(8):3940-9. PubMed PMID: 21289114.

46. Sigal A, Kim JT, Balazs AB, Dekel E, Mayo A, Milo R, et al. Cell-to-cell spread of HIV permits ongoing replication despite antiretroviral therapy. Nature. 2011;477(7362):95-8. PubMed PMID: 21849975.

47. Yoder A, Guo J, Yu D, Cui Z, Zhang XE, Wu Y. Effects of Microtubule Modulators on HIV-1 Infection of Transformed and Resting CD4 T Cells. J Virol. 2011;85(6):3020-4. PubMed PMID: 21209111.

48. Guo J, Wang W, Yu D, Wu Y. Spinoculation triggers dynamic actin and cofilin activity facilitating HIV-1 infection of transformed and resting CD4 T cells. J Virol. 2011;85(19):9824-33. PubMed PMID: 21795326.

49. Boullé M, Müller TG, Dähling S, Ganga Y, Jackson L, Mahamed D, et al. HIV Cell-to-Cell Spread Results in Earlier Onset of Viral Gene Expression by Multiple Infections per Cell. PLOS Pathogens. 2016;12(11):e1005964. doi: 10.1371/journal.ppat.1005964.

50. Fu Y, He S, Waheed AA, Dabbagh D, Zhou Z, Trinite B, et al. PSGL-1 restricts HIV-1 infectivity by blocking virus particle attachment to target cells. Proc Natl Acad Sci U S A. 2020;117(17):9537–45. doi: 10.1073/pnas.1916054117. PubMed PMID: 32273392; PubMed Central PMCID: PMCPMC7196789.

51. Wei X, Decker JM, Liu H, Zhang Z, Arani RB, Kilby JM, et al. Emergence of resistant human immunodeficiency virus type 1 in patients receiving fusion inhibitor (T-20) monotherapy. Antimicrob Agents Chemother. 2002;46(6):1896-905. PubMed PMID: 12019106.

52. Wu Y, Marsh JW. Selective transcription and modulation of resting T cell activity by preintegrated HIV DNA. Science. 2001;293(5534):1503–6.

53. Wu Y, Marsh JW. Early transcription from nonintegrated DNA in human immunodeficiency virus infection. J Virol. 2003;77(19):10376-82. PubMed PMID: 12970422.

54. Meltzer B, Dabbagh D, Guo J, Kashanchi F, Tyagi M, Wu Y. Tat controls transcriptional persistence of unintegrated HIV genome in primary human macrophages. Virology. 2018;518:241–52. doi: 10.1016/j.virol.2018.03.006. PubMed PMID: 29549786; PubMed Central PMCID: PMCPMC6021179.

55. Korichneva IL, Grigorian G, Krasnikova TL, Rudchenko SA, Tkachuk VA. Interleukin-2-and phytohemagglutinin-activated proliferation of human T-lymphocytes is accompanied by stimulation of phosphoinositide turnover. Biochim Biophys Acta. 1989;1014(2):173–7. doi: 10.1016/0167-4889(89)90030-x. PubMed PMID: 2554974.

56. McLinden RJ, Labranche CC, Chenine AL, Polonis VR, Eller MA, Wieczorek L, et al. Detection of HIV-1 neutralizing antibodies in a human CD4(+)/CXCR4(+)/CCR5(+) T-lymphoblastoid cell assay system. PLoS One. 2013;8(11):e77756. doi: 10.1371/journal.pone.0077756. PubMed PMID: 24312168; PubMed Central PMCID: PMC3842913.

57. Sarzotti-Kelsoe M, Daniell X, Todd CA, Bilska M, Martelli A, LaBranche C, et al. Optimization and validation of a neutralizing antibody assay for HIV-1 in A3R5 cells. J Immunol Methods. 2014;409:147–60. doi: 10.1016/j.jim.2014.02.013. PubMed PMID: 24607608; PubMed Central PMCID: PMC4138262.

58. Huskens D, Van Laethem K, Vermeire K, Balzarini J, Schols D. Resistance of HIV-1 to the broadly HIV-1-neutralizing, anti-carbohydrate antibody 2G12. Virology. 2007;360(2):294-304. PubMed PMID: 17123566.

59. Liu Y, Fu Y, Wang Q, Li M, Zhou Z, Dabbagh D, et al. Proteomic profiling of HIV-1 infection of human CD4(+) T cells identifies PSGL-1 as an HIV restriction factor. Nat Microbiol. 2019;4(5):813–25. doi: 10.1038/s41564-019-0372-2. PubMed PMID: 30833724.

60. Murakami T, Carmona N, Ono A. Virion-incorporated PSGL-1 and CD43 inhibit both cell-free infection and transinfection of HIV-1 by preventing virus-cell binding. Proc Natl Acad Sci U S A. 2020;117(14):8055–63. doi: 10.1073/pnas.1916055117. PubMed PMID: 32193343; PubMed Central PMCID: PMCPMC7148576.

61. Sheehy AM, Gaddis NC, Choi JD, Malim MH. Isolation of a human gene that inhibits HIV-1 infection and is suppressed by the viral Vif protein. Nature. 2002;418(6898):646-50. PubMed PMID: 12167863.

62. Usami Y, Wu Y, Gottlinger HG. SERINC3 and SERINC5 restrict HIV-1 infectivity and are counteracted by Nef. Nature. 2015;526(7572):218–23. doi: 10.1038/nature15400. PubMed PMID: 26416733; PubMed Central PMCID: PMCPMC4600458.

63. Rosa A, Chande A, Ziglio S, De Sanctis V, Bertorelli R, Goh SL, et al. HIV-1 Nef promotes infection by excluding SERINC5 from virion incorporation. Nature. 2015;526(7572):212–7. doi: 10.1038/nature15399. PubMed PMID: 26416734; PubMed Central PMCID: PMCPMC4861059.

64. Hammarskjöld ML, Heimer J, Hammarskjöld B, Sangwan I, Albert L, Rekosh D. Regulation of human immunodeficiency virus env expression by the rev gene product. Journal of Virology. 1989;63(5):1959–66. doi: doi:10.1128/jvi.63.5.1959-1966.1989.

65. Nabel G, Baltimore D. An inducible transcription factor activates expression of human immunodeficiency virus in T cells. Nature. 1987;326(6114):711-3. PubMed PMID: 3031512.

66. Shaw J-P, Utz PJ, Durand DB, Toole JJ, Emmel EA, Crabtree GR. Identification of a Putative Regulator of Early T Cell Activation Genes. Science. 1988;241(4862):202–5. doi: doi:10.1126/science.3260404.

67. Harrich D, Garcia J, Wu F, Mitsuyasu R, Gonazalez J, Gaynor R. Role of SP1-binding domains in in vivo transcriptional regulation of the human immunodeficiency virus type 1 long terminal repeat. Journal of Virology. 1989;63(6):2585–91. doi: doi:10.1128/jvi.63.6.2585-2591.1989.

68. Garcia JA, Harrich D, Soultanakis E, Wu F, Mitsuyasu R, Gaynor RB. Human immunodeficiency virus type 1 LTR TATA and TAR region sequences required for transcriptional regulation. The EMBO Journal. 1989;8(3):765–78. doi: 10.1002/j.1460-2075.1989.tb03437.x.

69. Spandidos DA, Yiagnisis M, Pintzas A. Human immunodeficiency virus long terminal repeat responds to transformation by the mutant T24 H-ras1 oncogene and it contains multiple AP-1 binding TPA-inducible consensus sequence elements. Anticancer Res. 1989;9(2):383-6. PubMed PMID: 2665635.

70. Hamer DH, Leder P. Splicing and the formation of stable RNA. Cell. 1979;18(4):1299–302. doi: 10.1016/0092-8674(79)90240-X.

71. Romfo CM, Ann Wise J. Both the polypyrimidine tract and the 3′ splice site function prior to the first step of splicing in fission yeast. Nucleic Acids Research. 1997;25(22):4658–65. doi: 10.1093/nar/25.22.4658.

72. Richardson MW, Jadlowsky J, Didigu CA, Doms RW, Riley JL. Kruppel-like Factor 2 Modulates CCR5 Expression and Susceptibility to HIV-1 Infection. The Journal of Immunology. 2012;189(8):3815–21. doi: 10.4049/jimmunol.1201431.

73. Hawley T, Spear M, Guo J, Wu Y. Inhibition of HIV replication in vitro by clinical immunosuppressants and chemotherapeutic agents. Cell Biosci. 2013;3(1):22. Epub 2013/05/16. doi: 2045-3701-3-22[pii]10.1186/2045-3701-3-22. PubMed PMID: 23672887; PubMed Central PMCID: PMC3680316.

